# CRISPR/Cas9 mediated gene editing in non-model nematode *Panagrolaimus* sp. PS1159

**DOI:** 10.1101/2022.09.26.509268

**Authors:** Viktoria Hellekes, Denise Claus, Johanna Seiler, Felix Illner, Philipp H. Schiffer, Michael Kroiher

## Abstract

The phylum Nematoda harbors a huge diversity of species in a broad range of ecosystems and habitats. Nematodes share a largely conserved Bauplan but major differences have been found in early developmental processes. The development of the nematode model organism *Caenorhabditis elegans* has been studied in great detail for decades. These efforts have provided the community with a large number of protocols and methods. Unfortunately, many of these tools are not easily applicable in non-*Caenorhabditis* nematodes. In recent years it has become clear that many crucial genes in the *C. elegans* developmental toolkit are absent in other nematode species. It is thus necessary to study the developmental program of other nematode species in detail to understand evolutionary conservation and novelty in the phylum.

*Panagrolaimus* sp. PS1159 is a non-parasitic nematode exhibiting parthenogenetic reproduction and we are establishing the species to comparatively study evolution, biodiversity and alternative reproduction and survival strategies. Here, we demonstrate the first successful application of the CRISPR/Cas9 system for genome editing in *Panagrolaimus* sp. PS1159 and its closely related hermaphroditic species *Propanagrolaimus* sp. JU765 with both the non-homologous end joining and the homology-directed repair mechanism. Using microinjections and modifying published protocols from *C. elegans and P. pacificus* we induced mutations in the orthologue of *unc-22*, which resulted in a visible uncoordinated twitching phenotype. We also compared the HDR efficiency following the delivery of different repair templates. This work will expand the applicability for a wide range of non-model nematodes from across the tree and facilitate functional analysis into the evolution of parthenogenesis, changes in the developmental program of Nematoda, and cryptobiosis.

## 1 Introduction

Understanding the molecular machinery controlling cellular fate during development of an organism is a major goal in developmental biology. Functional analysis of genes and how they are regulated aids to understand how, for example, the body shape is encoded in genomes (Levine and Davidson, 2005). The phylum Nematoda comprises of an estimated 1 million species (Lambshead and Boucher, 2003) occupying a broad range of habitats including terrestrial, marine, and aquatic environments (Holterman et al., 2019). Many species have adopted a parasitic lifestyle, or have evolved survival strategies to deal with extreme cold and desiccation (Blaxter and Koutsovoulos, 2015; Shannon et al., 2005). Throughout the Nematoda, different modes of reproduction are realized. For example, parthenogenesis, a form of asexual reproduction, occurs frequently. Despite their different biotopes and lifestyles, all nematodes share a morphologically similarly structured and largely conserved Bauplan (Poinar, 2011). In contrast to this conserved outcome of development, variations are found in early development on the cellular level with divergent cleavage patterns (Dolinski et al., 2001; Schulze and Schierenberg, 2009). This variability of embryogenesis shows that the patterns and structures observed in a single species, such as the derived nematode model *Caenorhabditis elegans*, do not allow a generalization across the phylum. We are particularly interested in parthenogenetic nematodes, which initiate development without male input (as present in *C. elegans* where sperm entry defines embryonic axis-formation (Goldstein and Hird, 1996)). To understand parthenogenesis as a trait it is necessary to establish new study systems.

On the genetic level, many aspects of the molecular machinery for development have been studied and are understood in the nematode model system. However, it has recently become clear that many developmentally crucial *C. elegans* genes are absent in many other nematode species. This includes for example sex determination, axis formation and endo-mesoderm specific genes (Kraus et al., 2017; Schiffer et al., 2013, 2018). It is thus necessary to study functions of genes and their regulations throughout the diversity of the phylum. Some other nematode species have been studied to greater detail, for example *Pristionchus pacificus*, a species from the same clade as *C. elegans* (Figure 1), but we lack approachable and informative systems from across the tree. Parthenogenesis, for example, is not present in any of the more than 50 *Caenorhabditis* species named to date, nor in species close to *Pristionchus*.

**Figure 1:**
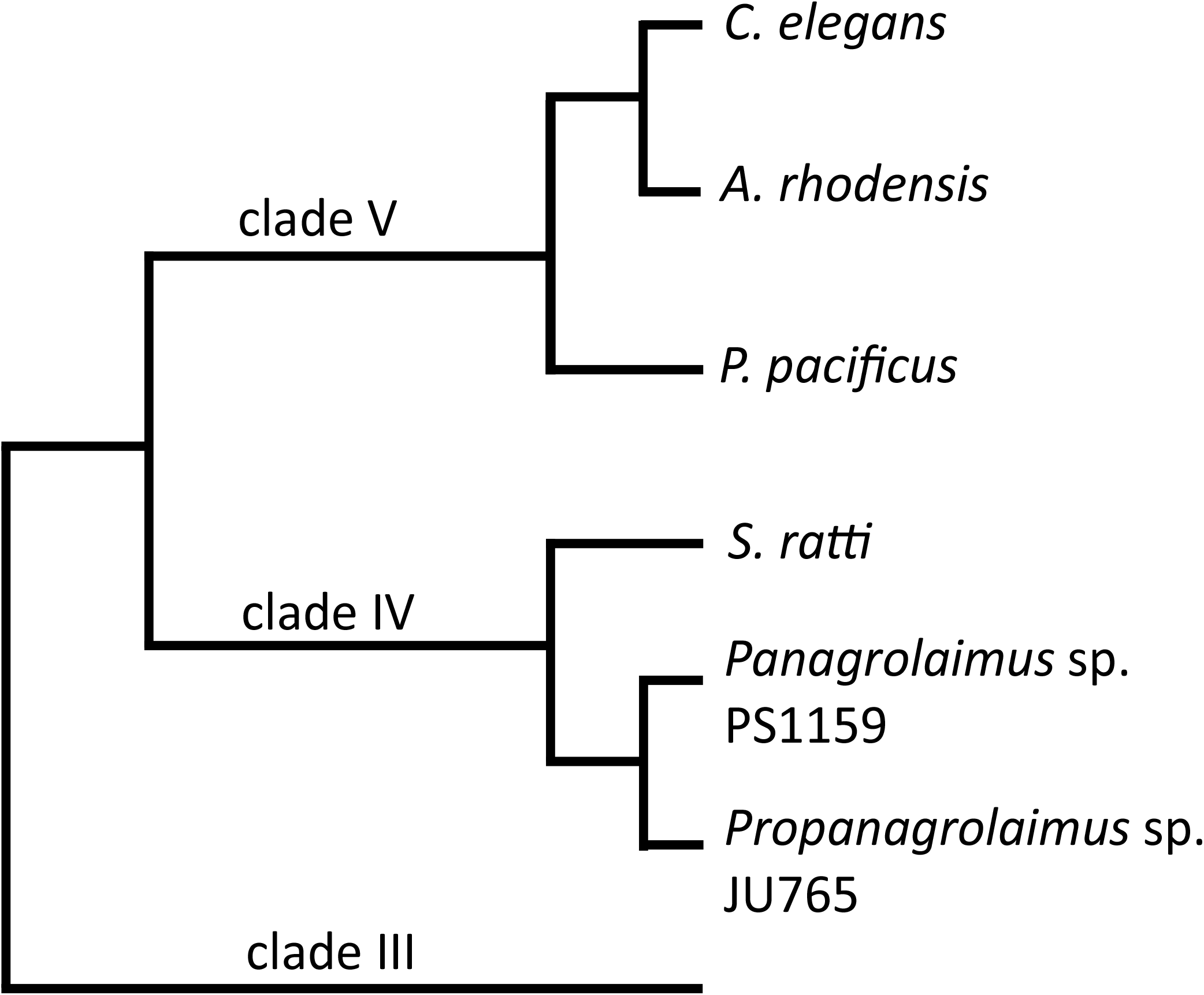
A cladogram showing the position of the two main species in this study, *Panagrolaimus* sp. PS1159 and *Propanagrolaimus* sp. JU765, in relation to the nematode model organisms *C. elegans* and *P. pacificus*, as well as *A. rhodensis* and *S. ratti*, species with reportedly successful use of the CRISPR system (modified from Blaxter and Koutsovoulos, 2015; Tandonnet et al., 2018)

In nematodes many species can easily be cultured in the laboratory, and the availability of more high-quality genomes through recent advances in sequencing technology is enabling us to define precise targets for gene knock-out studies. Unfortunately, many methods and protocols established in the model organisms do not work out-of-the-box in non-*Caenorhabditis* nematodes and organisms with less background on the genetic level. For example, RNAi has been used in a range of nematodes (Adams et al., 2019; Dulovic and Streit, 2019; Ratnappan et al., 2016; Shannon et al., 2008), but very often shows issues in efficiency and delivery (Félix, 2008), as nematodes are often insensitive to this method and important genes of the pathway are missing (Koutsovoulos, pers. comment). In our hands, RNAi by feeding did not result in reliable phenotypes in different parthenogenetic roundworms, e.g., *Diploscapter coronatus, Acrobeloides nanus* and *Panagrolaimus* sp. PS1159. The CRISPR technology promises to be more reliable, delivering consistent results across a broader range of species and thus opening new opportunities for functional analysis in many nematodes.

CRISPR/Cas9 is a powerful experimental tool for gene-editing (Wang et al., 2016). The simple design, its high efficiency, and low cost have made it the go-to-method in non-model organisms in recent years (Dickinson et al., 2020). The most important and widely adapted CRISPR/Cas application for gene editing is the type II system, which relies on one protein only (Makarova et al., 2011). The *Streptococcus pyogenes* Cas9 requires two small RNAs to build a ribonucleoprotein complex (RNP): the CRISPR RNA (crRNA) with a 20 bp guide sequence and its downstream protospacer-adjacent motif (PAM; NGG) that determines the target, and the universal trans-activating CRISPR RNA (tracrRNA). The crRNA binds to the DNA target by base-pair binding. Cas9 recognizes the PAM sequence, leading to a cleavage of both strands 3 nucleotides away from the PAM (Doudna and Charpentier, 2014), followed by cellular repair mechanisms. The Non-homologous End Joining (NHEJ) repairs the double-strand break (DSB) by ligating the ends. This mechanism sometimes results in the loss of nucleotides (Davis and Chen, 2013), a frame shift, premature stop and therefore creating knock-out mutations. The homology directed repair (HDR) is a more accurate system, in which user-provided DNA templates with homology arms serve as a guide to precisely repair the DSB (Mali et al., 2013; Paix et al., 2017) generating knock-in mutations. Cas9 can be delivered to the target DNA in various ways, for example via expression plasmids, or as a purified protein (Lino et al., 2018). It was shown that direct injection of the RNP complex into the adult gonad is more efficient and less time consuming than injection of plasmids (Paix et al., 2015). Over the past years, CRISPR/Cas9 was successfully applied in *C. elegans*. But the system has been proven to be useful also in non-model nematodes, such as *C. briggsae* (Culp et al., 2020), *P. pacificus* (Witte et al., 2015), the human- and rat-parasitic worms *S. stercoralis* and *S. ratti* (Gang et al., 2017), *Auanema* (Adams et al., 2019) and *O. tipulae* (Vargas-Velazquez et al., 2019).

Here we report the successful and reproducible use of microinjection and CRISPR/Cas9 in two new nematode strains from clade IV (Figure 1), the parthenogenetic strain *Panagrolaimus* sp. PS1159 and the hermaphroditic species *Propanagrolaimus* sp. JU765.

We also provide an optimization of previously published protocols using modified microinjection needles, different target regions as well as temperature and concentration adjustment during the RNP complex assembly. Additionally, we compare the HDR efficiency following the delivery of different repair templates.

## 2 Results

### 2.1 Microinjection in *Panagrolaimus* sp. PS1159

In nematodes CRISPR/Cas9 components are delivered through injection into the adult gonad. We found the cuticle in *Panagrolaimus* to be more robust and harder to breach than in *C. elegans*. At the same time the gonad appeared to be less tolerant to the injury resulting by the needles used for *C. elegans*. Thus, for successful microinjection in *Panagrolaimus* sp. PS1159 we adapted the *C. elegans* microinjection protocol (Evans, 2006) accordingly (see supplementary video V1).

*Panagrolaimus* worms were also less likely to stick on the 2% agarose pads. We therefore used higher concentrations of approximately 2.5% agarose pads for good adhesion of the worms. To prepare injection needles using a puller (Sutter Instruments, P-2000), we use a 1-lined so called “bee stinger” program (Oesterle, 2018) as a basis for a 3-lined program (table 1), producing sharper needles with a short taper. With these modifications we were able to easily inject *C. elegans, P. pacificus*, PS1159, and JU765 with a high survival rate.

**Table 1:** Needle puller P-2000 (A) One-line program “bee stinger needles” B) newly written 3-line program for our microinjection needles

| Program | Line | Heat | Filament | Velocity | Delay | Pull |
| --- | --- | --- | --- | --- | --- | --- |
| A | 1 | 350 | 4 | 40 | 200 | 0 |
|  | 1 | 350 | 4 | 40 | 200 | 0 |
| B | 2 | 350 | 4 | 40 | 200 | 0 |
|  | 3 | 450 | 4 | 60 | 130 | 100/150 |

### 2.2 Gene knock-out of *unc-22* in PS1159

We targeted the PS1159 orthologue of the *C. elegans unc-22* gene, which was previously successfully mutagenized by CRISPR/Cas9 in *C. elegans* (Kim et al., 2014), and the non-model nematodes *S. stercoralis*, and *S. ratti* (Gang et al., 2017). *unc-22* encodes Twitchin, a large protein, which is involved in the regulation of muscle contraction (Moerman et al., 1988). Mutation of *unc-22* in *C. elegans* results in motility defects and a twitching phenotype, which is easily detectable under the dissecting microscope.

#### 2.2.1 Identification of the PS1159 *unc-22* ortholog

The Twitchin protein (UNC-22, isoform a) of *C. elegans* was downloaded from www.wormbase.org (WormBase ID CE33017) and blasted against the PS1159 genome (PRJEB32708) using the TBLASTN tool. A potential orthologue with 62.9% sequence similarity was selected (“PS1159_v2.g4256”). Reciprocal BLASTN against the *C. elegans* genome confirmed the *C. elegans unc-22* gene as the best hit indicating the gene found is an *unc-22* orthologue. Target guide RNA sequences, preferably following the form G(N)G(NGG) were searched for in the exonic regions of the *unc-22* orthologue in our PS1159 genome assembly (Schiffer et al., 2019) using CRISPOR (Figure 2A; Concordet and Haeussler, 2018). Guide sequences with a GG motif at the 3’ end have been observed to have a higher cleavage efficiency in *C. elegans* (Farboud and Meyer, 2015). BLASTN of the chosen guide sequence revealed no significant off-target sequences in the entire genome. Two pairs of repair templates with homology arms of eighter 36 nt on both sides or 96 and 92 nt flanking the predicted Cas9 cleavage site with a built-in mutation and restriction enzyme *Xba*I site were produced (Figure 2A).

**Figure 2:**
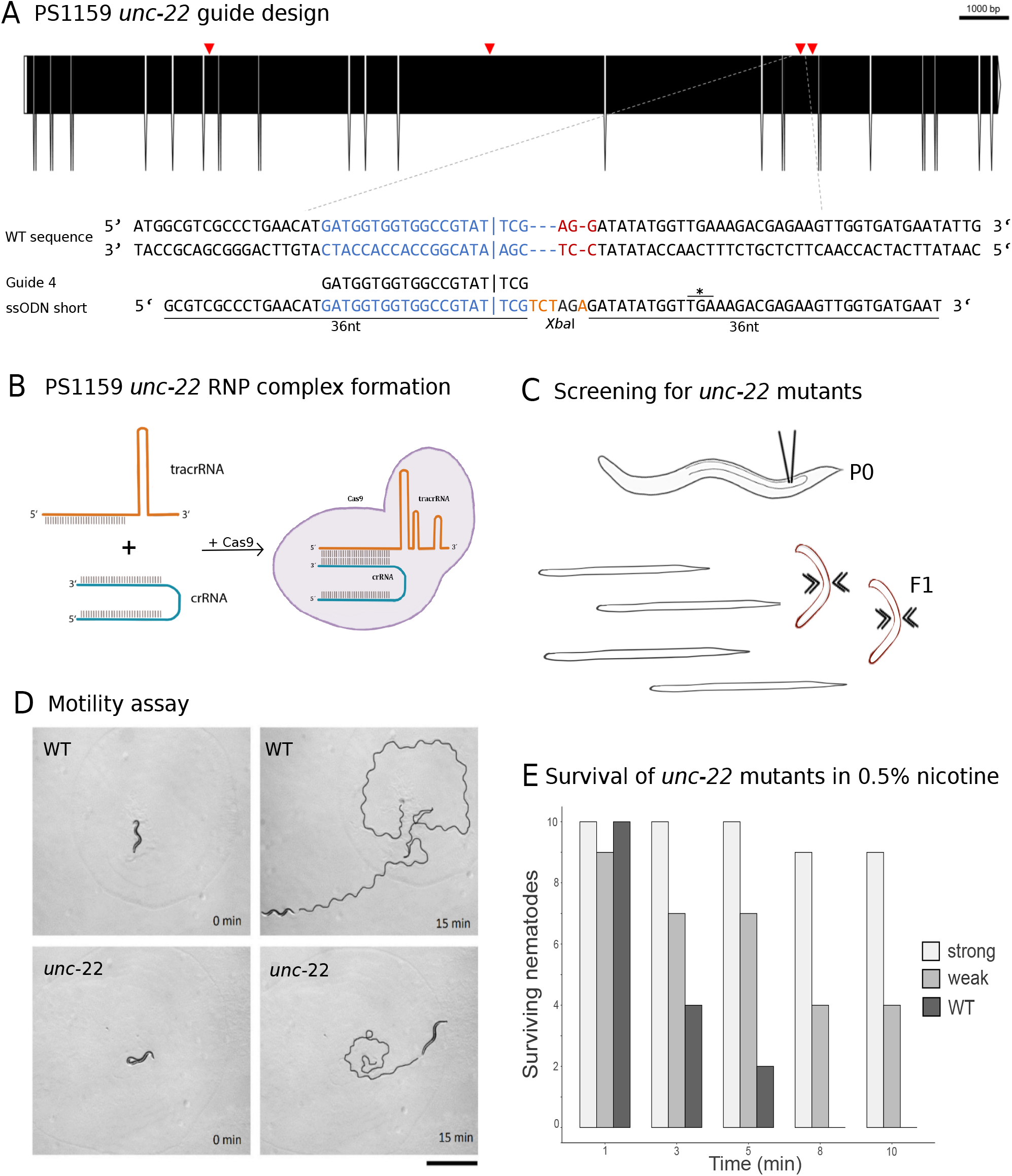
**(A)** CRISPR target site for *Panagrolaimus* sp. PS1159 *unc-22*. Predicted gene structure. Black boxes = exons. Red arrows = CRISPR target regions. Zoom into part of exon 15 coding strand with target region (blue) and PAM site (red). Vertical bar indicates the expected Cas9 cleavage site. Example of guide sequence and single-stranded oligodeoxynucleotide (ssODN) homology arms (36 nt) flanking the cleavage site on both sides and insertion of a *Xba*I cleavage site (orange). Asterisk shows in frame stop codon after successful incorporation of desired bases. Gene figure was designed by using the Exon-Intron Graphic Maker (version 4, http://wormweb.org/exonintron) **(B)** Ribonucleoprotein complex assembly. Cas9 protein, crRNA and tracrRNA are mixed and incubated **(C)** RNP complexes are introduced into the syncytial part of the gonad of an adult worm (P0) by microinjection. Screen for twitching phenotype in F1 generation **(D)** Motility assay: Observation of motility of wild-type worms vs. strong twitching F1 in the time course of 15 min (n=10); black line shows movement; scale bar = 1 mm. **(E)** Survival assay: Wild type worms and F1 progeny after injection with a phenotype (strong and weak) are screened in 0.5% nicotine for a total of 10 minutes for survival (n=10 for each)

#### 2.2.2 CRISPR-Cas9 targeting of PS1159 *unc-22* causes an uncoordinated phenotype

CRISPR/Cas9 components were assembled and introduced into the syncytial part of the gonad of PS1159 following the protocol of Paix et al, 2017. After injections, all worms were recovered on a single agar plate. After 12 days (generation time is 8-10 days in *Panagrolaimus*; Schiffer et al., 2019), 16 F1 animals showing reduced crawling activity and a twitching phenotype in water were transferred into single drops of *Plectus* nematode growth medium (PNGG; supplementary Table S1). We were able to isolate 4 different twitchers with varying intensity of their twitching behavior, most likely indicating weak (C-shaped bending to the left and right side with twitching observable in head and tail region in liquid) and strong (no bending to both sides, strong twitching through whole body) phenotypes (Supplementary video V1). This phenotype was visible and consistent in all of the progenies of future generations. The motility on agar plates of strong twitchers was monitored and compared to wild-type worms. The movement of twitching worms appeared uncoordinated. In a comparative assay of 10 wild type worms versus 10 strong twitchers, wild type nematodes appeared to have moved a longer distance after 15 minutes, indicating reduced motility in mutants (Figure 2D).

In *C. elegans*, wild-type worms stop moving and eventually die under exposure of nicotinic acetylcholine receptor agonists, such as levamisole or nicotine. *C. elegans unc-22* mutants are resistant to these poisons and twitching is enhanced by exposure (Lewis et al., 1980; Moerman et al., 1988). We tested exposure of PS1159 wild-type and putative CRISPR-induced *unc-22* mutants to both, levamisole (1mM, not shown) and 0.5% nicotine. 10 wild-type animals and 10 of each twitching phenotypes were analyzed (Figure 2C, E; supplementary video V1). After 10 minutes all worms were rescued in M9 buffer. No wild-type animal, 9 strong and 5 weak twitchers survived the procedure, suggesting that like in *C. elegans* wild-type PS1159 are not resistant to this poison and that the resistance of worms with different twitching phenotypes seems unequal. For all future experiments the progeny of injected worms was screened in water or, if confirmation was needed, additionally in 0.5% nicotine for 2-3 minutes, ensuring not to lose possible mutants. We also tested 1300 wild-type PS1159 for spontaneous twitching behavior in water and 1% nicotine for 10 minutes to analyze if this phenotype could frequently occur in a wild-type population, but could not observe any.

### 2.3 Development of an optimal cultivation method after injections for PS1159

The transfer of single worms per plate, as has been done in *C. elegans* and *P. pacificus*, was not optimal for *Panagrolaimus* sp. PS1159, because the worms frequently crawled out of the agar to the walls of the plates and dried out. To find the optimal way of cultivating the worms after microinjections, different approaches were taken. Injected worms were transferred either (1) into single PNGG drops on 12-well plates in a moist chamber individually, (2) all individuals onto a single agar plate or (3) onto single plates with each worm covered by a slice of agar. Although surviving the transfer into single PNGG drops, worms could not be screened properly because of strong bacterial growth in the drops. When worms were placed on a plate altogether, they remained on the plate but analysis of F1s from individual P0s was impossible. In the latter approach, almost all worms remained under the agar pad and laid eggs. Occasionally some worms still left and dried out, but the advantage of this method outweigh the loss of single individuals. However, since the reproduction cycle of PS1159 is much longer, than in *C. elegans* and *P. pacificus*, mothers were kept at 25 °C for at least 5-7 days on the plates, allowing them to lay more eggs.

### 2.4 Modification of the CRISPR mix preparation increases efficiency

During the first experiments, we were only able to isolate a few worms with a twitching phenotype and injection results were thus unsatisfactory and not consistent. After the development of the optimal cultivation strategy and the possibility to screen progeny of individual injected P0s, we aimed to adapt the protocol of the CRISPR injection mix to yield a higher efficiency.

#### 2.4.1 Using guide RNAs targeting different regions in the gene shows only minor variations, but using a modified crRNA increases the efficiency dramatically

To examine if the genomic region of the double strand break affects the editing success, we used 4 targets in different exons (Figure 2A; Table 4). The total outcome of progeny with the desired phenotype after injection of 60 µM crRNA varied between 1.14% and 2.63% (Table 2). When using a modified crRNA from IDT (Alt-R CRISPR-Cas9 crRNA XT), which contains additional chemical modifications providing an increased stability the efficiency rose dramatically to up to 16%.

**Table 2:**
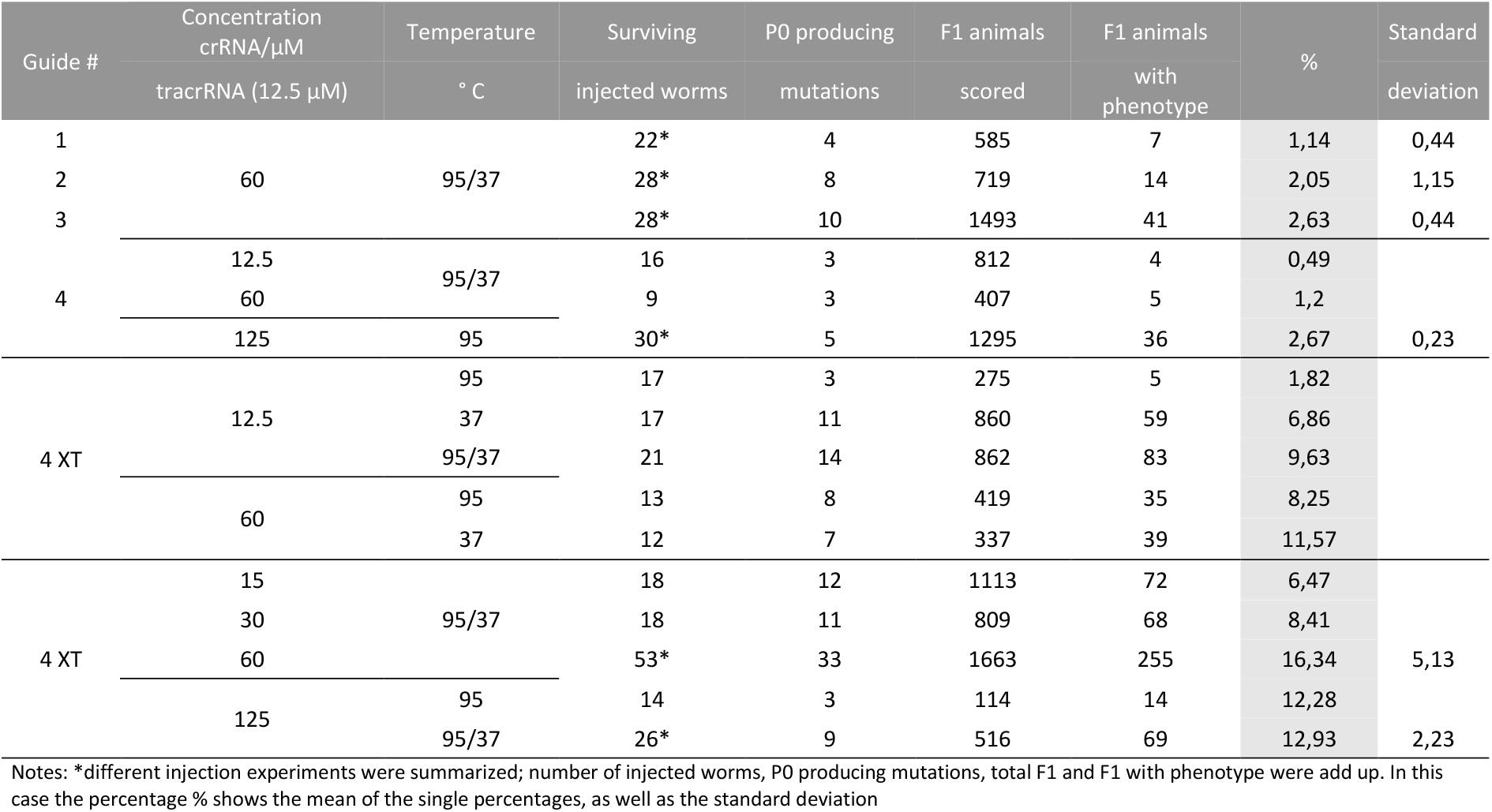
Summary of CRISPR/Cas9 experiments targeting the PS1159 *unc-22* gene

#### 2.4.2 Changes of temperature optimizes complex formation

In *Pristionchus pacificus* the CRISPR mix protocol differs from the one used in *C. elegans* (Paix et al., 2017) by an additional pre-annealing step of the tracrRNA and crRNA (James Lightfoot, pers. comment). We tested if the incubation temperature has an impact on the complex formation in PS1159. We therefore injected the complex with the modified crRNA with 60 µM concentration. When pre-annealing the crRNA and tracrRNA at 95 °C for 5 minutes (Hiraga et al., 2021) at the beginning we yielded 8.25% efficiency. When assembling the mix without the annealing step at 95 °C, but with an incubation step at 37 °C for 10 minutes after adding the Cas9 we reached to 11.57% of twitching progeny. When using both steps, the percentage of F1s with the phenotype increased up to 16.34 (Table 2).

#### 2.4.3 Guide RNA concentration determines effect-size of gene editing

We then asked how the concentration of the crRNA affected the mutation frequency. We used concentrations of 12.5 µM, 15 µM, 30 µM, 60 µM and 125 µM and discovered that by increasing up to 60 µM improved the efficiency rate but did not further increase when going up to 125 µM (Table 2). After these findings we decided that our future experiments would be performed with that optimal concentration of 60 µM.

#### 2.4.4 The optimal time-window for screening is during the first 48 hours after injection

After establishing the enhanced protocol for the injection mix, we further investigated the optimal time window for observing CRISPR induced knock-outs in PS1159. After microinjections, P0s were isolated onto single agar plates with a slice of agar and transferred every 24 hours for a total of 6 days (Figure 3A). We found that in the first 24 hours the average percentage of mutated progeny was 25% and then dropped to 8.5% after 48 hours (Figure 3B). After 72 hours, there was still an average of 7.5% and then 0% to 1.2% in the next 3 days. From these results we decided that the preferred time window to screen for mutations after injections, is 48 hours post injection after considering the feasibility and simplicity of maintenance of the progeny and achieving mutations arising from a higher number of P0s.

**Figure 3:**
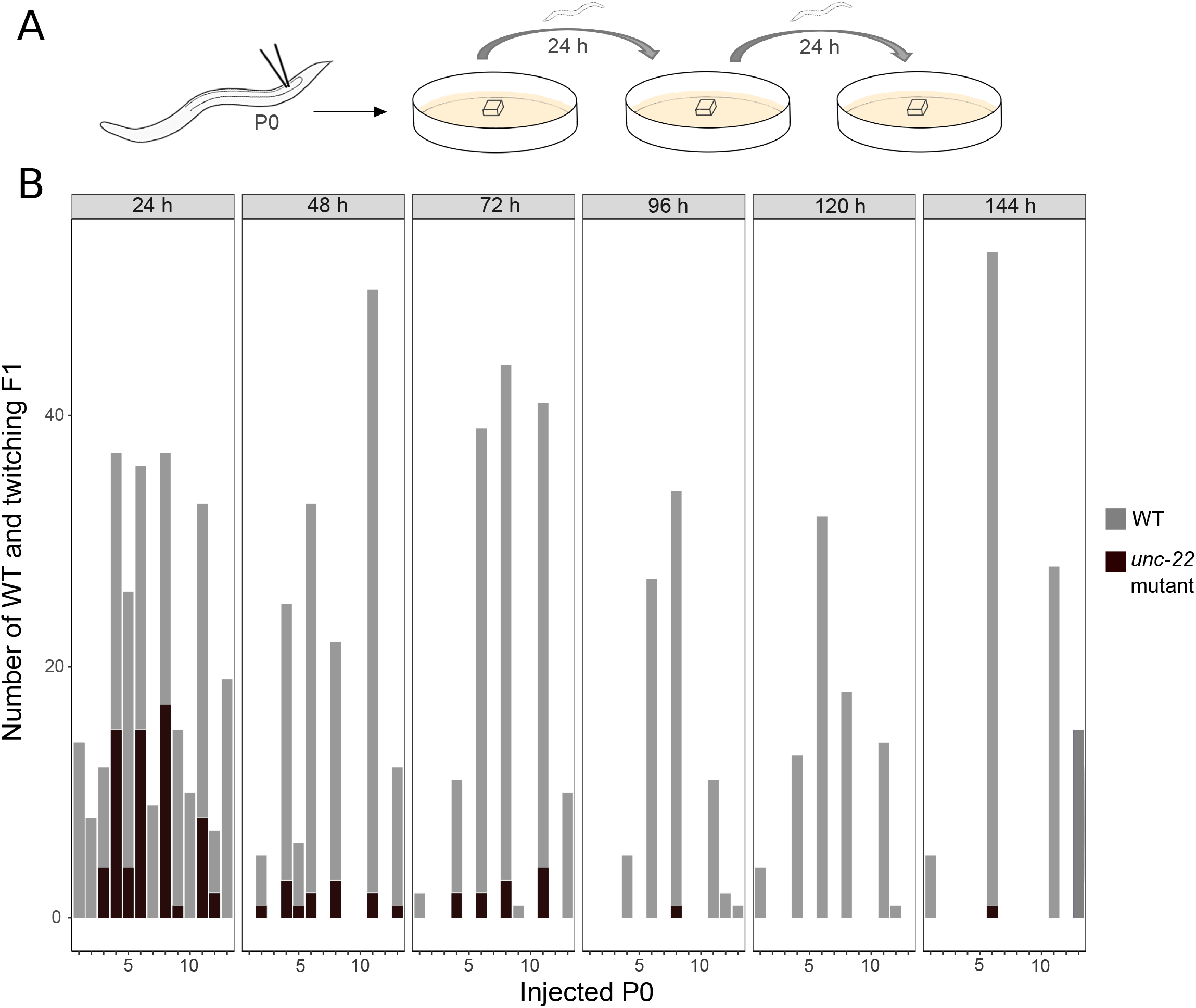
**(A)** Scheme of experimental workflow. Microinjection of adult P0 into gonad. Every 24 h individual worms are transferred onto fresh agar plates covered by a slice of agar **(B)** Number of wild-type (light grey) and twitching (dark grey) progeny per injected P0 in time intervals of 24 h for a total of 6 days

### 2.5 Sequencing verifies *unc-22* mutation after double strand break within the target site

We aimed to molecularly characterize the mutations produced after knock-out and therefore amplified the target region using primers located 328 bp upstream and 454 bp upstream of the PAM and sequenced it after single worm lysis and PCR. Sanger sequencing of the PCR product confirmed different types of mutations near the cleavage site (Figure 4A). To quantify and identify the genotypes from the trace data, we used the ICE tool from Synthego (Conant et al., 2022; Figure 4B). To determine the exact location of mutation, we cloned and subsequent sequenced each homeolog (for lack of an established nomenclature we here refer to potential “alleles” as homeologs in the triploid hybrids). The results were in accordance with the prediction from the Synthego Analysis tool (Figure 4C).

**Figure 4:**
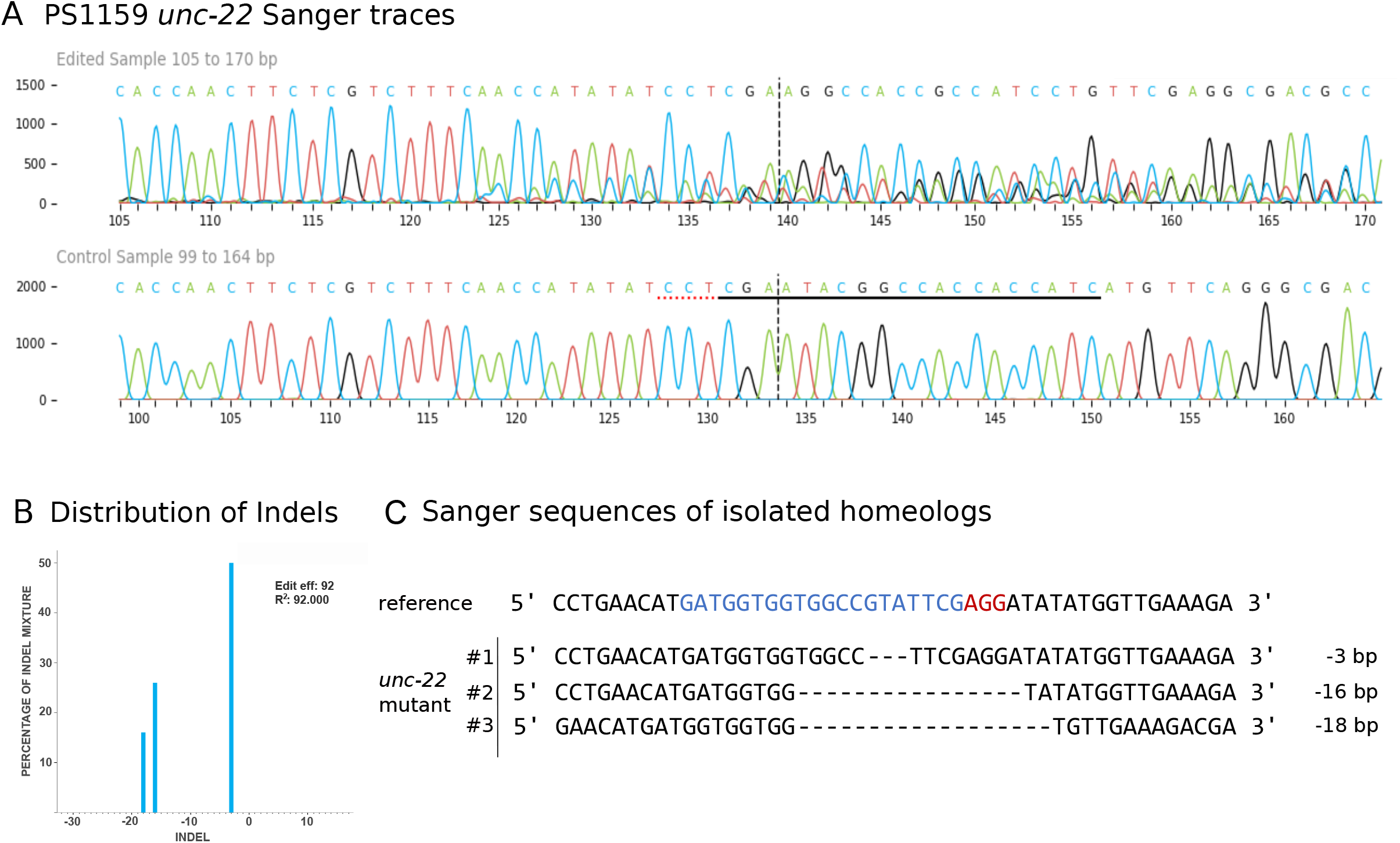
**(A)** *Panagrolaimus* sp. PS1159 *unc-22* Sanger Sequencing traces after PCR of edited and wild-type (control) in the region around the guide sequence using the Synthego Performance Analysis, ICE Analysis. 2019. v3.0. Synthego. Black underlined region shows the guide sequence. Red underline represents the PAM site. Vertical dotted line shows expected cut site **(B)** Predicted distribution and percentage of Indels in the entire population of genomes using ICE **(C)** Sanger sequencing reveals deletions in isolated homeologs #1-#3 after DNA cloning. Target region (blue) and PAM site (red).

In some cases, the amplification failed and only the WT sequences were observed even though a clear phenotype was visible, suggesting the deletion of at least one primer site. We attempted to localize the exact site of the deletion by increasing the PCR amplification region to up to 6000 bp and used different primer combinations. We found an amplification at 6000 bp (WT) and 3000 bp (supplementary Figure S1 A) of one mutant (T4) and at 3000 bp when using only reverse primers. We next sequenced this region and found a deletion around the target and forward primer site and an inversion of the reverse primer region (supplementary Figure S1 B).

### 2.6 Longer homology arms with modification increase efficiency

To simplify the detection of knock-in success we used homology arms with a built-in insert containing a recognition site for the restriction enzyme *Xba*I, which additionally delivers an in frame stop, thus allowing us to easily check for gene editing in the progeny with a phenotype via restriction digest (Figure 5A) after PCR. Restriction digest was successful and Sanger sequencing after DNA cloning confirmed the desired insertion in one homeolog (Figure 5B). We designed longer homology arms (97/92nt) with an additional incorporated phosphorothioate (PS) bond modification and observed the effect of knock-in efficiency when compared to the results of injections with 36 nt arms flanking the target site. We found that in 10% of twitching progeny the desired knock-in has been generated with short homology arms whereas using long modified arms with 97/92 nt resulted in a significantly higher outcome of knock-ins with 58% (n=30).

**Figure 5:**
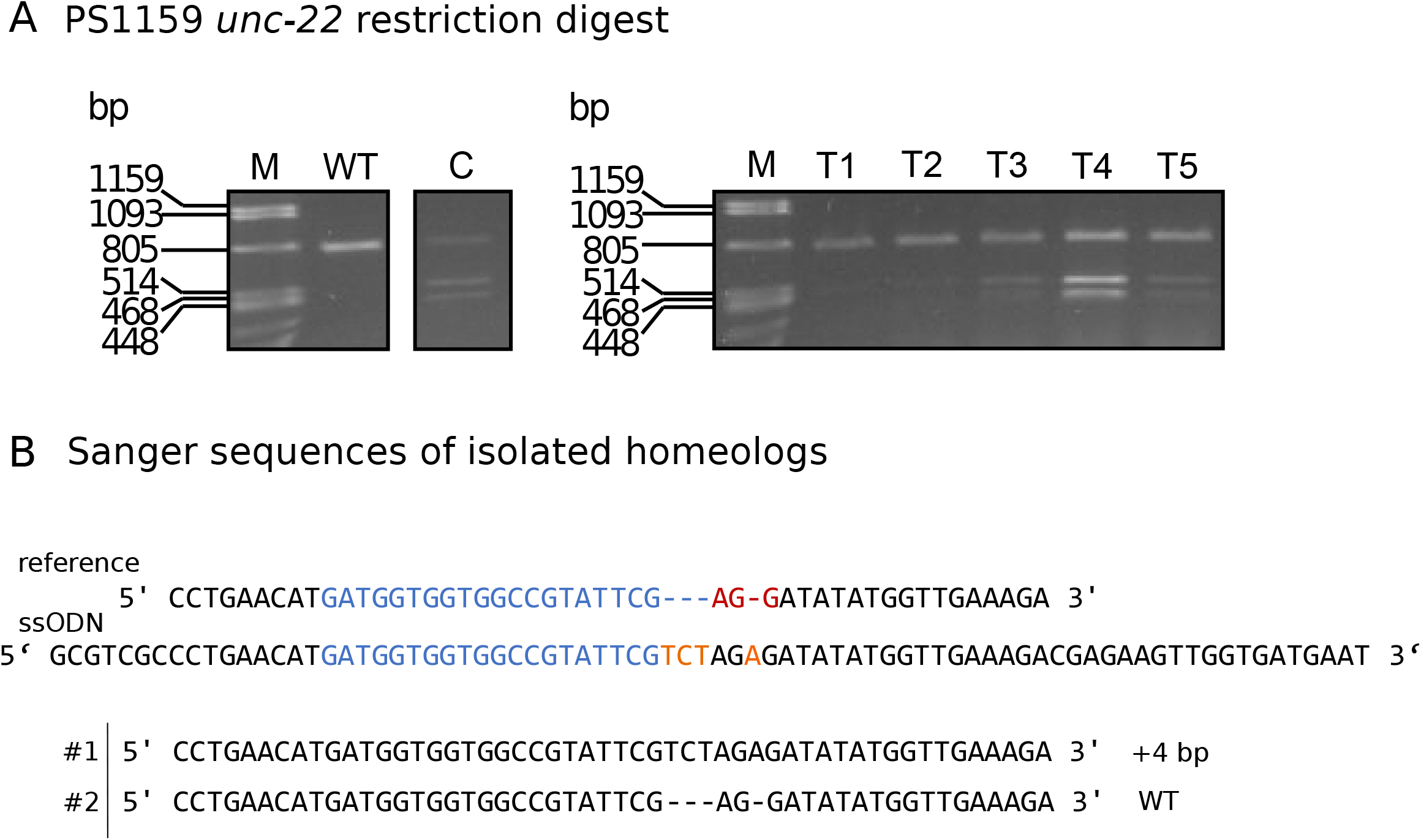
**(A)** *Panagrolaimus* sp. PS1159 *unc-22*: Representative examples of agarose gel electrophoresis after restriction digest with *Xba*I of knock-in experiments. Multiple band patterns of mutants with desired insertion. M=Marker Lambda *Pst*I digest. C=positive control of approved knock-in positive F1. T1 - T5= twitchers 1-5 **(B)** Sanger sequencing revealing insertion in one homeolog (#1) after DNA cloning of mutant after positive restriction digest. Wild-type sequence in homeolog #2. Target region (blue) and PAM site (red). *Xba*I cleavage site in dark yellow.

### 2.7 The same protocol can be applied in *Propanagrolaimus* sp. JU765

Following our improved protocol for PS1159, we asked whether we could transfer the injection mixture assembly and microinjection strategy to modify the genome of other non-model nematodes. We hence aimed to induce mutations in the *Propanagrolaimus* sp. JU765 orthologue of *unc-22*. After successful injections of 13 individuals, we screened the F1 progeny after 7 days and recovered 30.7% with a twitching phenotype (total F1=501) which could be observed in the next generations as well. To verify that the phenotype arose from a CRISPR induced knock-out, we amplified the target region with primers 437 bp upstream and 416 bp downstream of the PAM and performed a T7EI assay indicating a mutation within the region (Figure 6A). We identified the predicted position of indels with Sanger Sequencing and ICE analysis tool for 3 representatives (Figure 6B). This result supports that we provide an optimal protocol that might be useful in a variety of non-model species in the phylum Nematoda.

**Figure 6:**
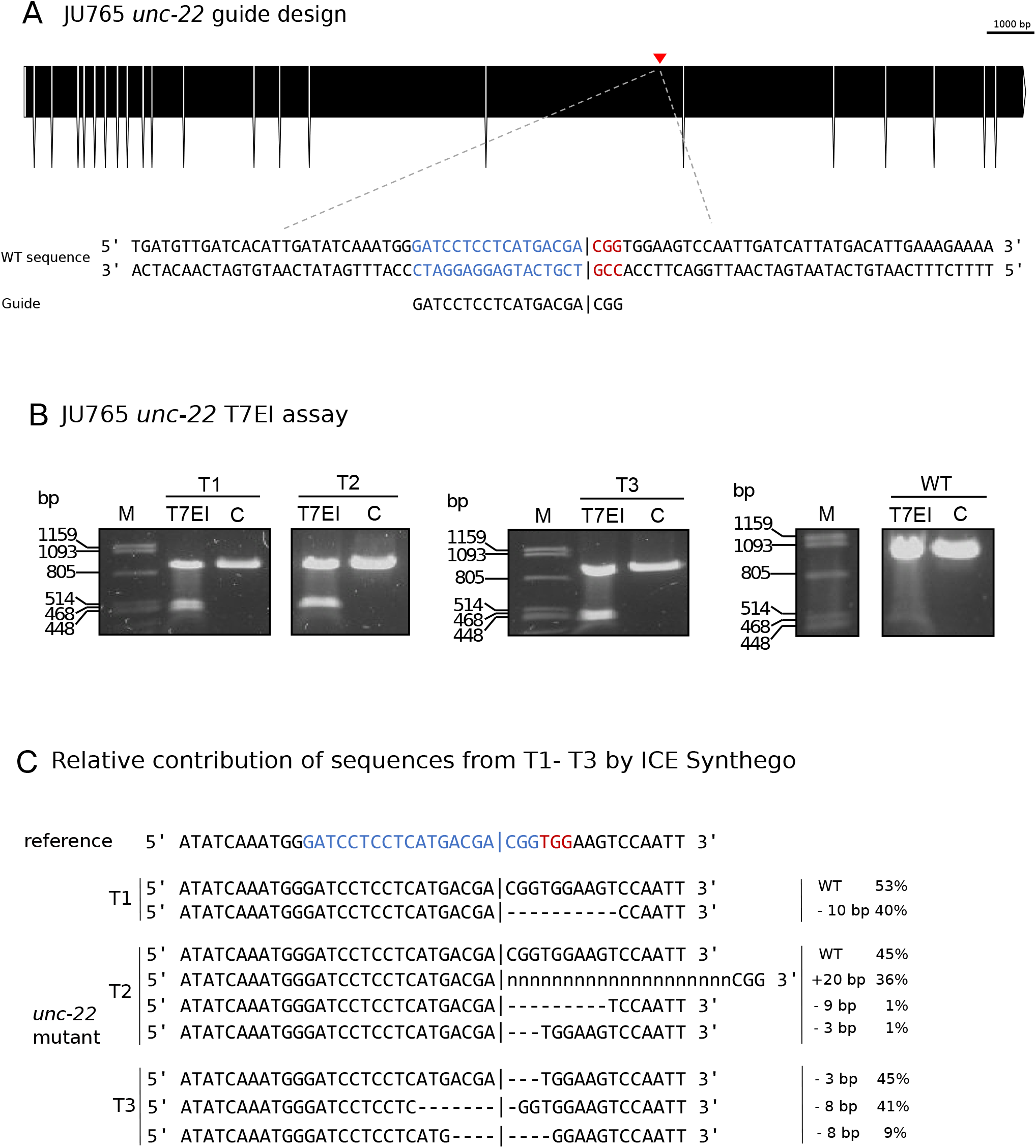
**(A)** CRISPR target site for *Propanagrolaimus* sp. JU765 *unc-22*. Predicted gene structure. Black boxes = exons. Red arrow = CRISPR target region. Zoom into part of exon 16 coding strand with target region (blue) and PAM site (red). Vertical bar indicates the expected Cas9 cleavage site. **(B)** Representative examples of 3 individuals T1-T3. Agarose gel electrophoresis of T7EI assay after knock-out experiments. Multiple band patterns detect indels. M=Marker Lambda *Pst*I digest. C=negative control. T1 - T3= twitchers 1-3 **(C)** Relative contribution of inferred sanger sequences in edited T1-T3 (PCR product) confirms indels in the target region. Target region (blue) and complementary PAM site (red). Vertical bar indicates the expected (actual) Cas9 cleavage site. Dashes indicate deletions and ‘n’ indicates insertions. The number of inserted (+) or deleted (-) nucleotides are indicated on the right with the proportion of that sequence inferred in the pool. Prediction produced using the Synthego Performance Analysis, ICE Analysis. 2019. v3.0. Synthego

## 3 Discussion

CRISPR/Cas9 has become an indispensable tool for precise gene editing in different eukaryotes, and has revolutionized genetics in the past years in many model organisms (Chang et al., 2013; Cho et al., 2013; Cong et al., 2013; Miki et al., 2021). In this study, we demonstrate the successful and efficient application of the CRISPR/Cas9 system in the nematodes *Panagrolaimus* sp. PS1159 and *Propanagrolaimus* sp. JU765 by generating heritable *unc-22* mutants. We further optimized the protocol for a higher cleavage efficiency.

Microinjection is a useful tool to introduce genetic material into an adult worm. In *C. elegans* this method has long been used for different approaches, including RNAi and CRISPR/Cas9 (Conte Jr. et al., 2015; Ghanta et al., 2021; Mello and Fire, 1995) and has successfully been used in some non-model nematodes as well (Adams et al., 2019; Ratnappan et al., 2016). Because of differences in the cuticle and gonad, the published protocols for *C. elegans* have not been working in our lab in PS1159. We therefore adapted existing protocols for the microinjection pads and needles. With these changes it was possible to inject not only PS1159, but also *C. elegans, P. pacificus*, and JU765 with high survival rates.

We generated knock-out mutations in the PS1159 and JU765 orthologues of the *C. elegans unc-22* gene which resulted in a visible uncoordinated twitching phenotype with reduced motility increased by nicotine exposure. This phenotype is consistent with what has been observed in *C. elegans* and *S. stercoralis*. To determine if the phenotype was caused by mutations in the desired target, we recovered and isolated single worms with a phenotype. We amplified, cloned, and sequenced the regions spanning the predicted sites. We were able to identify indels showing that the genotypes were consistent with the observed phenotypes. However, in some cases the amplification failed, and we only observed wild-type sequences which was in line with previous results from knock-out experiments in the nematodes *Strongyloides* and *Pristionchus* (Gang et al., 2017; Witte et al., 2015). We took a closer look and found a deletion of the guide sequence and an inversion of the reverse primer site, removing the forward primer site. It has been demonstrated in different organisms that repair of cleavage induced by Cas9 can lead to large deletions and genomic rearrangements using one or two single guide RNA’s (Kosicki et al., 2018; Shin et al., 2017) and should therefore be considered as a possible outcome. To create knock-in mutations of small insertions we provided single-stranded homology arms on both sites. We were able to achieve even higher knock-in efficiency when using longer homology arms with a protection of the donor DNA with phosphorothioate modification. This result is in accordance with previous findings regarding optimized HDR donors (Liang et al., 2017; Schubert et al., 2021).

We found that best results can be obtained when using a more stable, modified version of a crRNA with an additional pre-annealing step with the tracrRNA before incubation with the Cas9 protein. It has previously been reported that certain chemical alterations on terminal ends can lead to increased metabolic stability of guide RNA and subsequent higher CRISPR/Cas9 efficiency (Allen et al., 2021; Hendel et al., 2015). Increasing the concentration of the modified crRNA resulted in an increase from ∼1-2% to up to 16%. Moreover, we observed the editing efficiency in the timeframe of one week and count the frequency of mutations every 24 hours and our analysis indicates that the CRISPR effect in PS1159 performs at its best in the first 48 hours post injection, leading to even up to 25% cleavage efficiency. In *Pristionchus pacificus* it has been reported that most mutations arose in eggs laid in the first 9 hours after injections (Witte et al., 2015).

In summary, testing and adapting experimental conditions, such as needle size, temperature and concentration of the RNP complex components during preparation as well as using modified reagents is necessary for successfully applying microinjections and CRISPR/Cas9 in divergent nematode species. Our work will facilitate functional analysis into the evolution of parthenogenesis, changes in the developmental program of Nematoda, and cryptobiosis. The *unc-22* target might be a useful and easily identifiable co-marker for future functional analysis, in particular for large insertions of fluorescent markers in Panagrolaimidae, as it has been used in *C. elegans* (Arribere et al., 2014; Kim et al., 2014) routinely.

## 4 Materials and methods

### 4.1 Nematodes and maintenance

All experiments have been performed using the parthenogenetic *Panagrolaimus* sp. PS1159 and the hermaphroditic *Propanagrolaimus* sp. JU765, cultured on 55 mm diameter petri dishes at 20 °C with low-salt agar (Lahl et al., 2003) seeded with *Escherichia coli* strain OP50 as a food source (Brenner, 1974). After injections worms were kept at 25° C (20 °C for experiments with older protocol) either on single drops of 15 µl *Plectus* nematode growth medium (PNGG, supplemental Table S1; 30 µl OP50 in 1000 µl PNGG) on 12-well plates in a moist chamber, or on single agar plates covered with a slice of agar.

### 4.2 Selection of guide RNA targets

The *C. elegans* sequences of UNC-22 were downloaded from Wormbase (http://www.wormbase.org/db/get?name=CE33017;class=PROTEIN). TBLASTN and BLASTN tools (Altschul et al., 1990) were used to identify orthologues for PS1159 (PRJEB32708, Schiffer et al., 2019) and JU765 (PRJEB32708, Schiffer et al., 2019) in WormBase ParaSite (https://parasite.wormbase.org). For *Propanagrolaimus* sp. JU765 a potential orthologue with 63.5% similarity was selected (JU765_v2.g7216) We searched for potential guide RNAs and analyzed the putative off-targets using a web-based software (CRISPOR, http://crispor.tefor.net; Concordet and Haeussler, 2018). A list of guide RNA’s and ssODN’s can be found on table 3 and table 4.

**Table 3:**
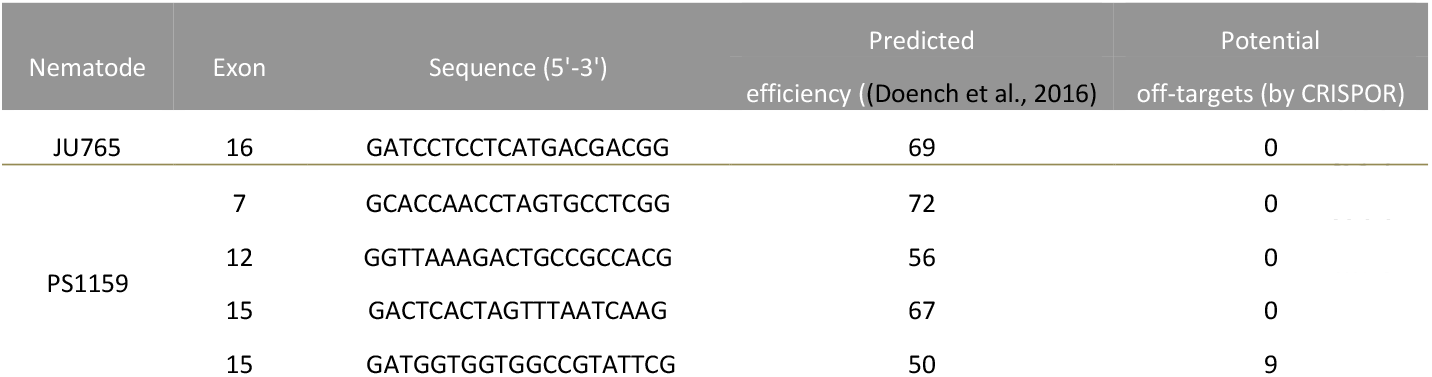
List of gRNAs used for this study

**Table 4:**
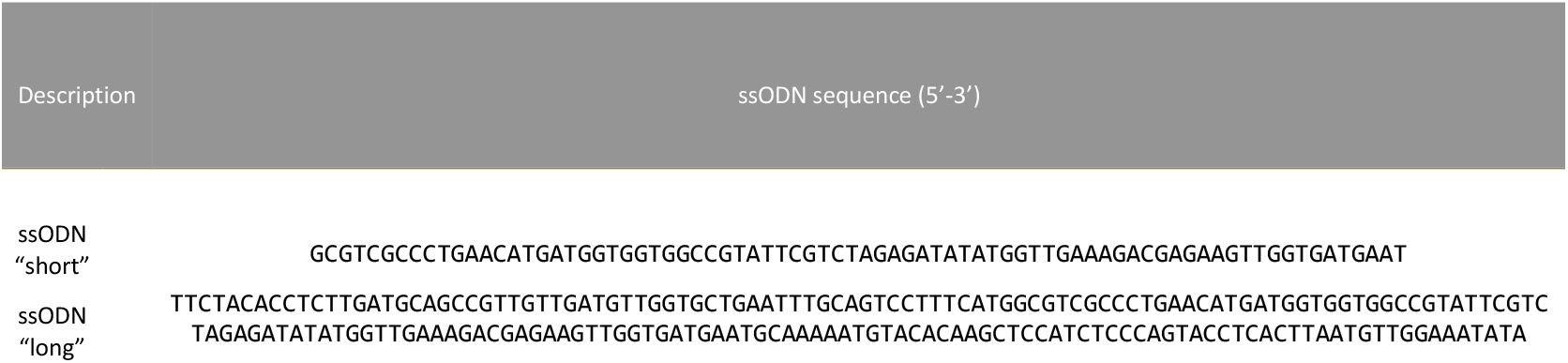
List of homology arms used in this study for *Panagrolaimus* sp. PS1159 *unc-22*

### 4.3 Preparation of HDR donor

For knock-in of small insertions using the homology directed repair (HDR), ssODN (single-stranded oligodeoxynucleotides) repair templates with 37 nt (“short”) and 97/92 nt (“long”) homology arms flanking the cleavage site with an inserted *Xba*I site followed by a stop codon were designed using Geneious Pro 5.5.6 (Kearse et al, 2012).

### 4.4 Preparation of injection mix

1.25 µl tracrRNA (100 µM, catalog# 1073191, IDT) and different amounts of crRNA according to the tested concentration of 12.5 µM, 30 µM, 60 µM and 125 µM (Alt-R® CRISPR-Cas9 crRNA (XT), 10 nmol, IDT) were mixed and incubated at 95 °C for 5 min following incubation at RT for 5 min. 0.5 µl Cas9 (10 mg/ml, catalog# 10811059, IDT) was added and incubated at 37 °C for 10 min for RNP (ribonucleoprotein) complex formation. 1 µl ssODN repair template (25 µM) was added and the mixture was filled up with TE buffer to 10 µl, centrifuged at 13.000 x g for 5 min and kept on ice for injections.

### 4.5 Preparation of injection needles, agar pads and microinjection

2.5% agarose pads were produced, and microinjections performed as described in Evans, 2006. PS1159 and JU765 are monodelphic and the syncytial region of the gonadal arm looks similar as in *C. elegans*. Borosilicate glass capillary (GB120F-10, 0.69×1.20×100 mm, Science products GmbH) were pulled in a needle puller (P-2000, Sutter instruments) and loaded with the final injection mixture. The needle was opened by gently tapping the tip against the edge of a broken piece of coverslip placed on an agar pad and covered with oil (Halocarbon oil 700, Sigma-Aldrich). An Inverted DIC microscope (Zeiss IM 35) equipped with standard Nomarski objectives (non-oil immersion, 6.3x and 40x) was used. A micromanipulator (Bachofer) with a needle holder with a fine mobility in the x, y and z axes (Piezo manipulator, PM10, Bachofer) was attached to the microscope and a pressure regulator (Pneumatic PicoPump PV 820) with a foot petal connection to a vacuum pump (Vacuubrand ME2C). Pressure was adjusted between 15 and 20 psi for injections. Worms were recovered in M9 buffer (Supplemental table S3), transferred to a single agar plate, and covered with a small piece of agar at 25 °C (video of microinjection steps in supplemental video V1). Phenol red dye was used for the purpose of the supplemental video V1. No dye was used for microinjection experiments.

### 4.6 Screening for knock-out mutations

#### 4.6.1 Single worm PCR and Sanger Sequencing

Individual worms were picked and transferred into tubes with 10 µl H_2_O or M9 buffer. Tubes were placed at –80 °C for at least 1 h. 10 µl 2x worm lysis buffer (supplemental Table S2) was added, incubated at 55 °C for 3 hours and heated at 95 °C for 10 minutes. We designed primers to amplify the genomic DNA fragment containing the cut site. PCR products were resolved on a 0.8% agarose gel. Sanger sequencing was performed through Eurofins Scientific SE. Sequences were analyzed using the ICE CRISPR analysis tool v3 (https://www.synthego.com; Conant et al., 2022) and Geneious Pro 5.5.6 (Kearse et al., 2012). List of primers used for this study can be found on table 5.

**Table 5:**
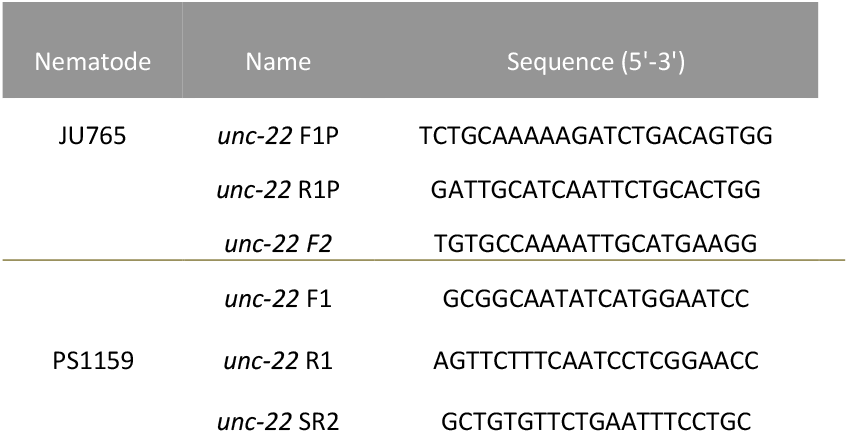
List of primers used for this study

#### 4.6.2 Motility assay

Observation of motility of wild-type worms vs. strong twitching F1 in the time course of 15 min (n=10); Single worms were transferred onto a drop of water in an agar plate. After the drop evaporated, the worms were allowed to acclimate for 1 minute and the sinusoidal waves of the movement were observed (Figure 2D).

#### 4.6.3 Survival assay

10 wild-type worms and 10 F1 progeny after injection with a phenotype (strong and weak) were screened in 0.5% nicotine for a total of 10 minutes for survival. Individual worms were first transferred into water and then nicotine. After 10 minutes worms were rescued and observed for survival in M9 buffer.

#### 4.6.4 T7 Endonuclease I assay

10 µl single worm PCR product was used for T7 Endonuclease I assay according to the protocol from New England Biolabs Inc. Samples were loaded onto a 1.4% agarose gel.

### 4.7 Screening for Knock-ins / HDR

#### 4.7.1 Restriction digest

10 µl single worm PCR products of F2 individuals were mixed with 1 µl Tango buffer, 1 µl *Xba*I enzyme and 13 µl H_2_O and incubated at 37 °C for 3 h. Samples were loaded in a 1.4% agarose gel.

### 4.8 Data analysis and editing

Gene structure diagrams were generated with the Exon-Intron Graphic Maker (Version 4, www.wormweb.org/exonintron). Plots were generated with R package ggplot2 (Wickham, 2016). Images and Videos were edited with InkScape 1.2, Adobe Illustrator CS6 v. 16.0.0 and Avidemux 2.8.0.

## 6 Author Contributions

VH conducted all major experiments, was involved in planning the experiments, analyzed the data, wrote the manuscript. DC conducted some experiments and contributed to figure design. JS conducted some experiments. FI contributed to one experiment. PS conceived the study, wrote the manuscript, and supervised VH and DC. MK conceived the study, planned the experiments, analyzed data, edited the manuscript, supervised VH, JS and FI and co-supervised DC.

## 8 Acknowledgments

We thank James Lightfoot for sharing RNP complex components and protocols for pilot experiments. This manuscript is dedicated to the memory of Einhard Schierenberg.

