## Supplementary files for "CRISPR/Cas9 mediated gene editing in non-model nematode *Panagrolaimus* sp. PS1159"

### *Supplementary Material*

#### 1 Supplementary Data

Supplementary video V1

#### 2 Supplementary Figures and Tables

##### 2.1 Supplementary Figures

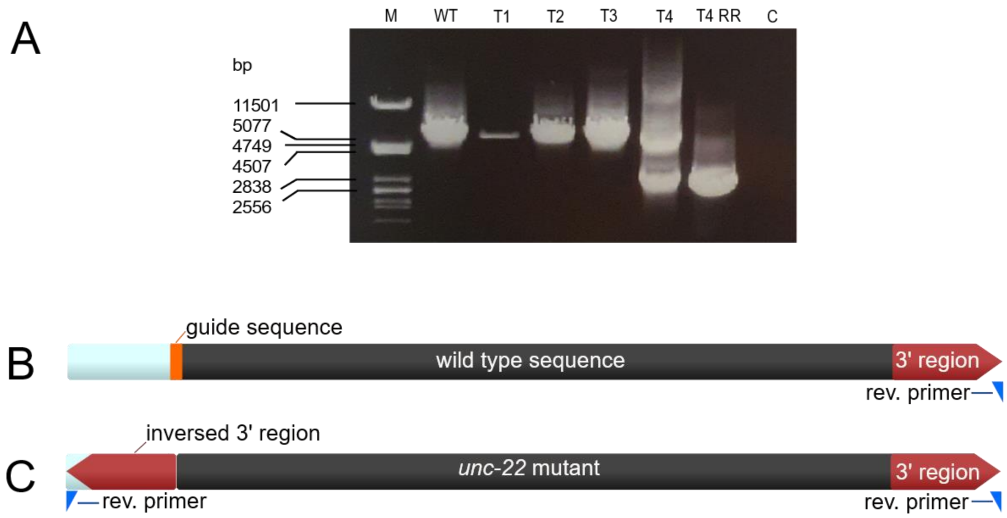


**Supplementary Figure S1. (A)** Agarose gel electrophoresis after PCR of 6000 bp region around the PS1159 *unc-22* target site with forward and reverse primers of mutant T1-T4. T4 RR= amplification of mutant T4 target region using reverse primers only. WT = wild type. C = Control. M = Marker Lambda *Pst*I digest **(B)** Schematic representation of wild type sequence around the *unc-22* target site **(C)** Schematic representation of *unc-22* mutant sequence (T4) with inversed 3' region and reverse primer site

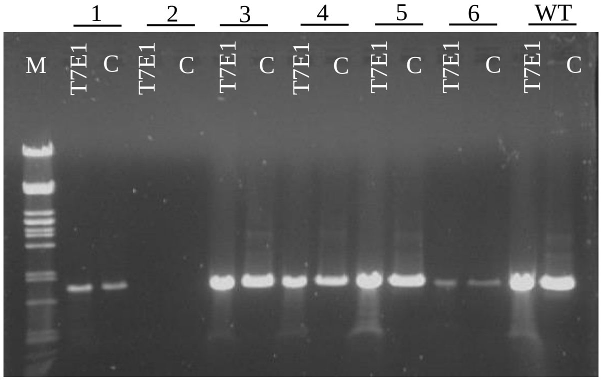


**Supplementary Figure S2.** Agarose gel electrophoresis after T7E1 assay of twitcher 1-6 (C=no T7 endonuclease, M=Marker Lambda *Pst*I)

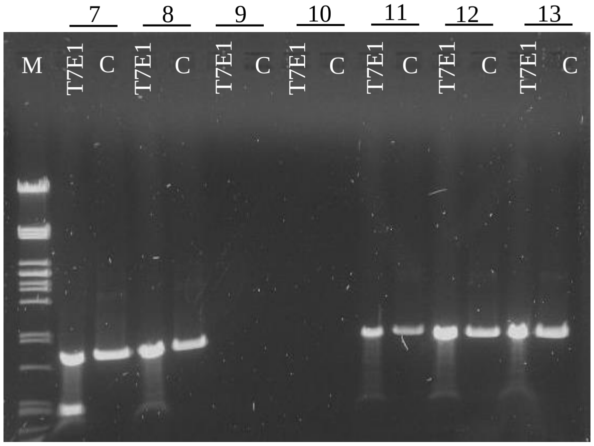


**Supplementary Figure S3.** Agarose gel electrophoresis after T7E1 assay of twitcher 7-13 (C=no T7 endonuclease, M=Marker Lambda *Pst*I); 7= T3 in figure 6

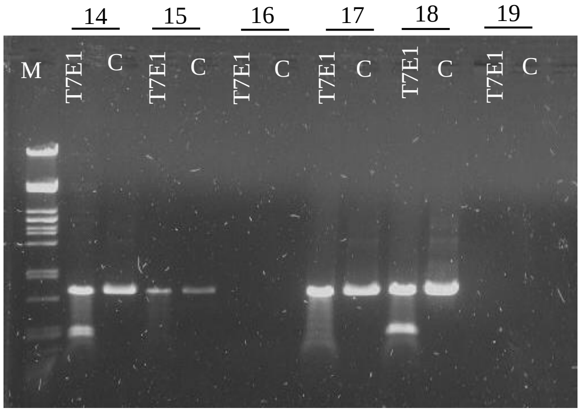


**Supplementary Figure S4.** Agarose gel electrophoresis after T7E1 assay of twitcher 14-20 (C=no T7 endonuclease, M=Marker Lambda *Pst*I); 14=T1, 18=T2 in Figure 6

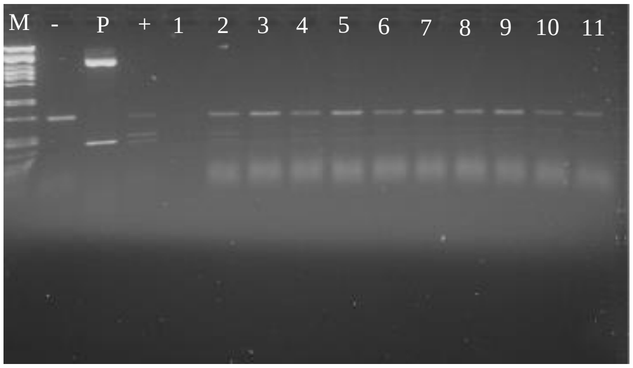


**Supplementary Figure S5.** Agarose gel electrophoresis after restriction digest of twitcher 1-11 (M=Marker Lambda *Pst*I, (-) negative control WT, (+) positive control, P=Plasmid with PstI restriction site)

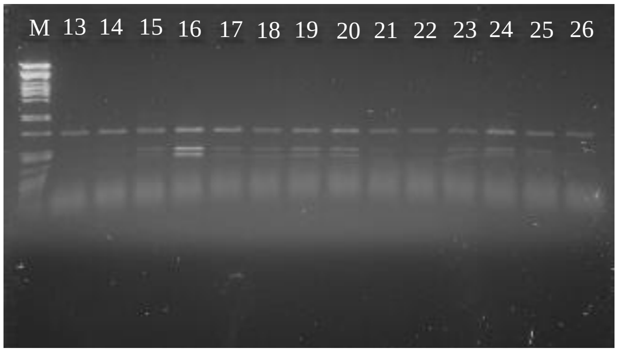


**Supplementary Figure S6.** Agarose gel electrophoresis after restriction digest of twitcher 13-26 (M=Marker Lambda *Pst*I, (-) negative control WT, (+) positive control, P=Plasmid with PstI restriction site); 13-17= T1-T5 in figure 5

#### 2.2 Supplementary tables

**Table S1:** PNGG growth medium

| **PNGG growth medium** |
|---|
| 0.3 g Gelrite (Roth # 0039.1) |
| Ad 200 ml H2O |
| Autoclave and cool to room temperature |
| 0.2 ml cholesterol (5mg/ml in 95% EtOH) |

**Table S2:** Worm lysis buffer

| **Worm lysis buffer** |
|---|
| 10 mM Tris pH 8.0 |
| 50 mM KCl |
| 2 mM MgCl2 |
| 0.5% NP40 |
| 0.5% Tween20 |
| 100 µg/ml Proteinase K |

**Table S3: M9 buffer**

| **M9 buffer** |
|---|
| 3 g $KH_2PO_4$ |
| 6 g $Na_2HPO_4$ |
| 0.5 g NaCl |
| 1 g $NH_4Cl$ |
| Bring to 1 L with $H_2O$. |
